# Cloning and functional characterization of *Komagataella phaffii* centromeres by a color-based plasmid stability assay

**DOI:** 10.1101/433417

**Authors:** Luiza Cesca Piva, Janice Lisboa De Marco, Lidia Maria Pepe de Moraes, Viviane Castelo Branco Reis, Fernando Araripe Gonçalves Torres

## Abstract

The yeast *Komagataella phaffii* is widely used as a microbial host for heterologous protein production. However, molecular tools for this yeast are basically restricted to a few integrative and replicative plasmids. Four sequences that have recently been proposed as the *K. phaffii* centromeres could be used to develop a new class of mitotically stable vectors. In this work we designed a color-based genetic assay to investigate genetic stability in *K. phaffii*. Plasmids bearing *K. phaffii* centromeres and the *ADE3* marker were evaluated in terms of mitotic stability in an *ade2/ade3* auxotrophic strain which allows plasmid screening through colony color. Plasmid copy number was verified through qPCR. Our results confirmed that the centromeric plasmids were maintained at low copy number as a result of typical chromosome-like segregation during cell division. These features, combined with high transformation efficiency and *in vivo* assembly possibilities, prompt these plasmids as a new addition to the *K. phaffii* genetic toolbox.

## Introduction

*Komagataella phaffii* is a methylotrophic yeast of great industrial importance which has been used for more than 30 years as a heterologous protein production platform [1]. Its genome was first published in 2009 and has since then been refined and thoroughly studied [2,3]. As a result, in addition to a protein factory, *K. phaffii* has also been widely considered as a platform for the production of chemicals, biopharmaceuticals, vitamins and other molecules. However, the construction and regulation of new pathways demand complex molecular biology tools which are not readily available for this yeast [4].

*K. phaffii* genetic manipulation traditionally involves the use of shuttle vectors assembled in *Escherichia coli* and subsequently integrated into the yeast’s genome [5]. Recent studies have described the development of a wide range of genetic parts for use in this yeast, as well as new methods of plasmid assembly and transformation [6]. An alternative to integrative strategies is the use of replicative plasmids, which are usually based on the well-known ARS1 sequence [1]. These plasmids may overcome some drawbacks such as genetic instability in multi-copy strains, non-specific genomic integration and different expression levels depending on the integration *locus* [7-9]. In addition, they present higher transformation efficiency when compared to integrative vectors and can be assembled by *in vivo* recombination, which eliminates the need for bacterial transformation [10,11]. However, replicative plasmids show low mitotic stability when compared to integrative vectors and few vector options are available for use [12]. Stability problems can be circumvented by the creation of centromeric plasmids, which may provide proper segregation during mitosis. Greater mitotic stability as well as low copy number allow stable and constant protein expression [13]. Centromeric plasmids can be constructed *in vivo,* allowing the assembly and cloning of large sequences including whole metabolic pathways and regulatory regions [14]. Therefore, the construction of such vectors would be of great value for *K. phaffii* strain development in the context of synthetic biology.

Centromeres are typically surrounded by large heterochromatin sections in most organisms [15]. Their structure ranges from simple “point” centromeres of only ∼125 bp in *Saccharomyces cerevisiae* to epigenetic, sequence-independent centromeres, such as those present in plants and animals. The reason for this phenomenon is that, for most eukaryotes, centromeres are maintained epigenetically and not genetically. Sequence homologies are rare in and between species, hindering the definition of a consensus sequence. In addition, some DNA regions can be centromeric or not depending on its function in previous cell cycles, which highlights the epigenetic nature of the centromere [16].

As for non-conventional yeasts, there are wide variations in centromere size and structure. *Candida glabrata* has centromeres that show some homology to the CDEI and CDEIII regions of *S. cerevisiae* while *Kuraishia capsulata* centromeres have 200-bp conserved sequences [17,18]. On the other hand, *Candida tropicalis, Schizosaccharomyces pombe* and *Candida albicans* have regional centromeres named after their sizes which range from 3 to 110 kb [19–21].

*K. phaffii* centromeres have recently been identified, bearing no sequence similarities to those of any other yeast [3]. Since centromere function relies strongly on its structure rather than on its sequence, a centromere-specific histone H3 variant (CSE4) was used in the search for centromeric regions in *K. phaffii*. A CSE4 homolog was identified in chromosome 2 and tagged with a fluorescence marker. The corresponding nuclear localization of the histone-DNA complex indicated a centromere pattern typical of budding yeasts [3]. Tridimensional conformation analysis followed the centromere clustering pattern observed in yeasts and narrowed down all four *K. phaffii* centromere locations to 20 kb windows [22].

Considering that a low transcription rate is typical of centromeric regions, RNA-seq analysis allowed researchers to pinpoint the putative centromeric locations for all four *K. phaffii* centromeres [3]. Similarly to *C. tropicalis* and *S.* pombe, *K. phaffii* centromeres are formed by inverted repeats. All four sequences have two inverted repeats of ∼2,5 kb, separated by a central segment of 800 to 1300 bp. Chromatin immunoprecipitation sequencing analysis showed that the CSE4 histone binds preferably to the central region, but also along the inverted repeats [23].

*K. phaffii* centromeric sequences contain early replication peaks with autonomously replicating sequences, characteristics that are also observed in centromeres of other yeasts [24,25]. According to recently published studies, there are native ARS sequences contained within centromeres 2, 3 and 4. These comprise regions within the inverted repeats, as well as unique adjacent sequences [23,26].

In order to expand the functional analysis of the *K. phaffii* centromeres we sought in this study to develop a genetic system based on an *ade2/ade3* auxotrophic strain and a replicative vector carrying the wild-type *ADE3*. Vectors carrying *K. phaffii* centromeres were used to assess plasmid copy number and mitotic stability.

## Results and Discussion

In yeasts, the adenine synthesis pathway is used as a tool for auxotrophic selection, gene copy number indicator and for plasmid stability analysis [27]. Many genes from this pathway have been deleted in *S. cerevisiae* in order to create auxotrophic strains, while in *K. phaffii* studies have only focused on *ADE1* and *ADE2* [28,29]. *K. phaffii* LA2, a strain mutant for *ADE2* [30], was used as a starting point for the construction of a strain that would allow plasmid stability verification. Deletion of *ADE2* results in cells that are auxotrophic for adenine and accumulate a red intermediate [27] while deletion of genes located upstream, such as *ADE1* or *ADE3*, should prevent the formation of such pigment [28]. As expected, *ADE3* deletion in the LA3 strain results in white colonies (Fig 1). The deletion of this gene in *S. cerevisiae* has regulatory effects in the histidine synthesis pathway [31]. Consequently, *ade2 ade3* strains are not only auxotrophic for adenine, but also for histidine. In order to verify if this phenotype is applicable to *K. phaffii*, we plated strains X-33, LA2 and LA3 on MD medium without supplementation, comparing growth and colony color to cells plated on MD medium with adenine and histidine (Fig 1). LA3 strain displayed the expected histidine auxotrophy phenotype, showing that the adenine-histidine pathways in *K. phaffii* and *S. cerevisiae* have common characteristics.

**Fig 1.**
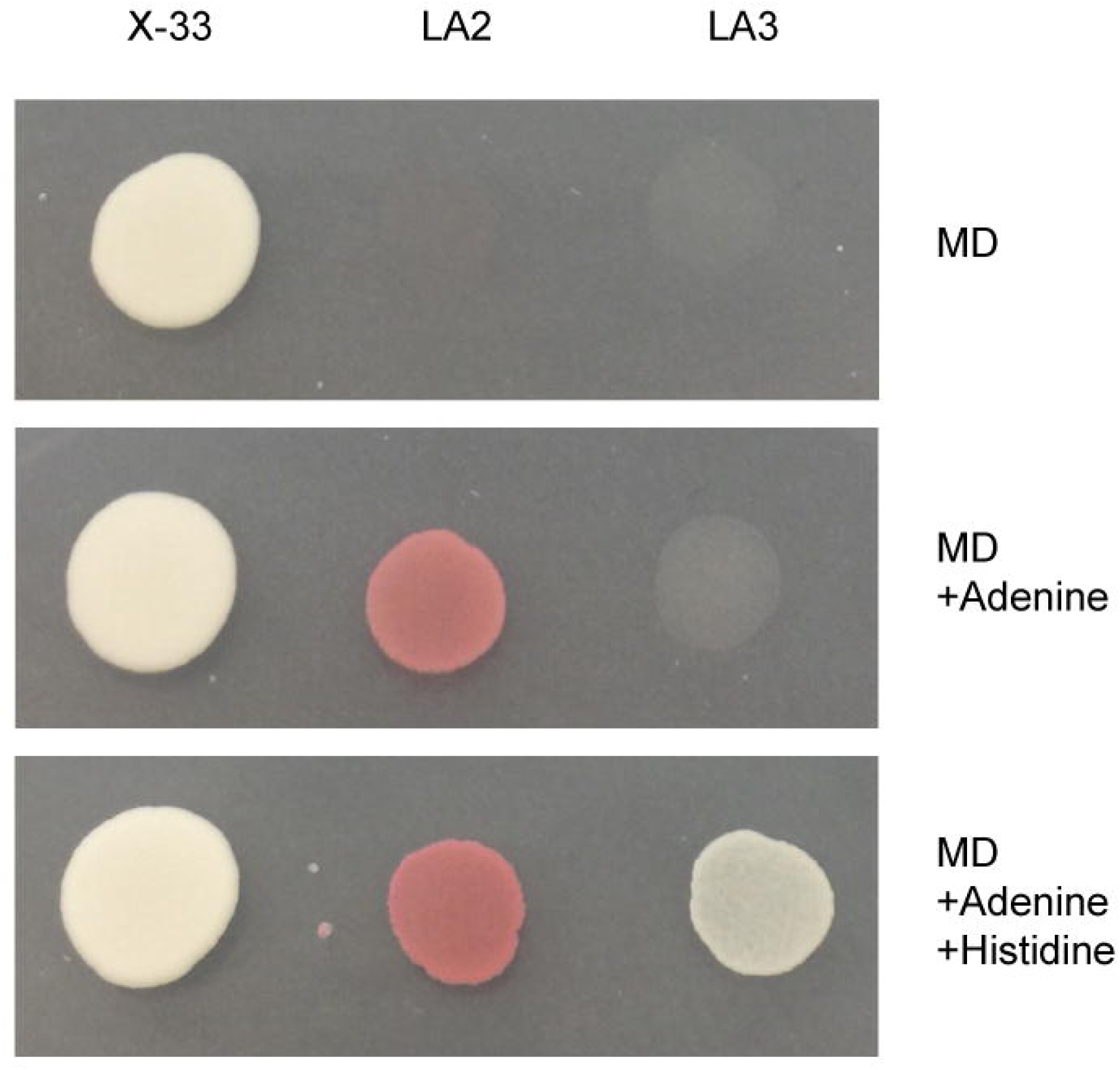
Strain phenotypic analysis on defined media. *K. phaffii* X-33 (wild-type), LA2 and LA3 cultures were spotted on MD medium with or without supplementation.

In order to assess plasmid stability, we first constructed plasmid pPICH-ADE3 bearing the *ADE3* gene (Fig 2). When transformed with pPICH-ADE3, LA3 cells should return to being red and any changes on colony color would allow a simple screening of plasmid loss [27]. Although adenine auxotrophy has been explored for other purposes in *K. phaffii* [29], this particular color-based system has not yet been used for measuring plasmid stability in this yeast.

**Fig 2.**
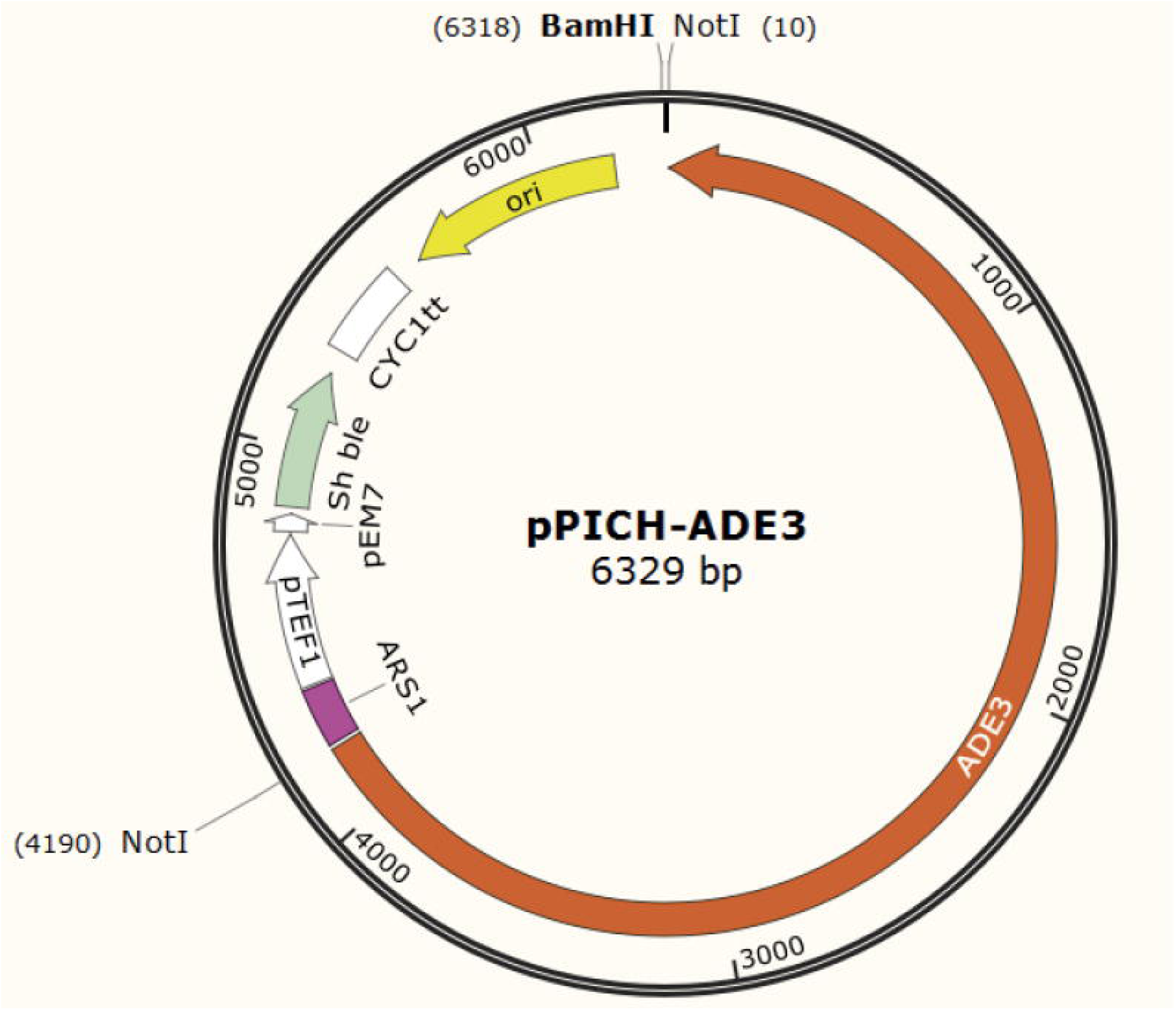
Map of vector pPICH-ADE3. ARS1 was used as the autonomously replicative sequence and *Sh ble* was used as the zeocin resistance marker. The *Not*I site was used for cloning the *K. phaffii ADE3* gene.

pPICH-ADE3 was used for cloning *K. phaffii* centromeres. Since it revealed extremely difficult to amplify entire centromeric regions, we designed a strategy to amplify centromeres in halves in order to reduce fragment size and to avoid primer annealing inside the inverted repeats (Fig 3). Amplified fragments exhibited in their ends overlapping regions that would allow recombination between each other and with vector pPICH-ADE3. Centromeric primer sequences were designed using *K. phaffii* GS115 genome sequence as reference [2]. The amplified regions corresponded to the following chromosomal coordinates: chromosome 1 position 1401429-1406917 (5488 bp); chromosome 2 position 1543739-1550657 (6918 bp); chromosome 3 position 2204800-2211493 (6693 bp) and chromosome 4 position 1703369-1709958 (6589 bp). Despite several attempts, we were unable to clone the centromere present in chromosome 3, which was excluded from our analysis. We speculate that its repetitive motifs, which were unlike those present in the other *K. phaffii* centromeres, rendered it extremely unstable in *E. coli*.

**Fig 3.**
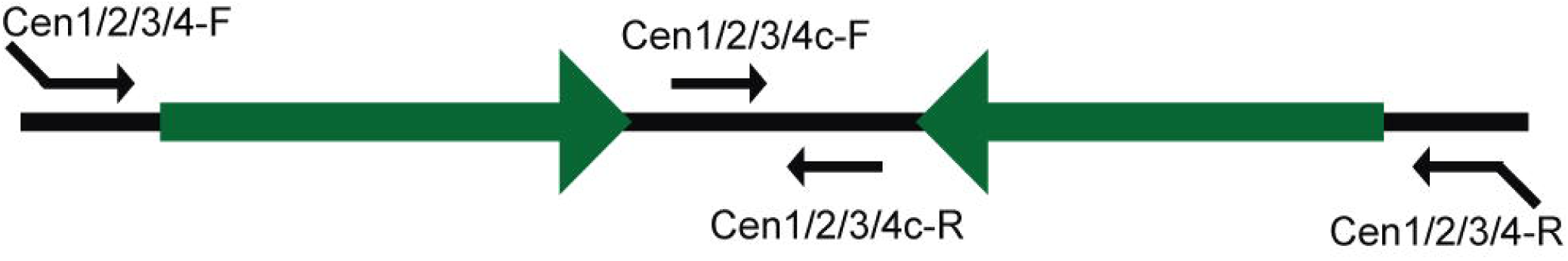
Strategy for *K. phaffii* centromere amplification. Schematic representation of a typical *K. phaffii* centromere. Inverted repeats are represented by green arrows. Primer annealing regions are shown by small arrows.

Centromeric sequences are known as early replication regions and according to recently published studies there are native ARS contained within centromeres 2, 3 and 4 [12, 23]. Fig 4 illustrates the positions of the ARS within and around these centromeres. In chromosome 2, ARS are located on coordinates 1543374-1543971 (597 bp) and 1549967-1551156 (1189 bp). These were partially amplified in this work, containing 232 and 690 bp, respectively. Regarding chromosome 3, there is an ARS located on coordinates 2204369-2205185 (816 bp) which was also partially amplified with 385 bp. Chromosome 4 has an ARS on coordinates 1703466-1704103 (637 bp), which was fully amplified, as well as a partially amplified one (840 bp) on coordinates 1709118-1710114 (996 bp total) [12].

**Fig 4.**
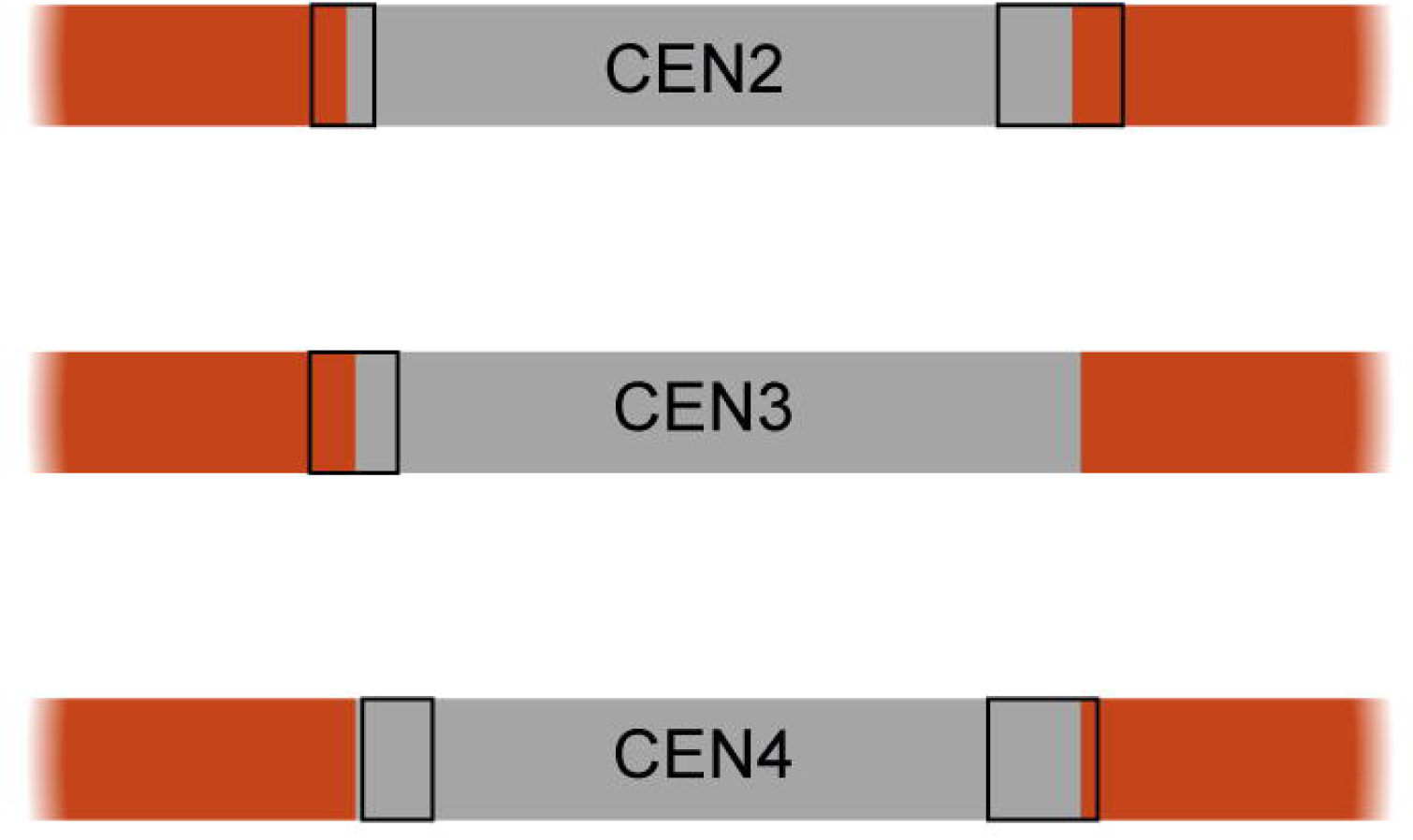
Relative positions of ARS sequences around centromeres 2, 3 and 4 of *K. phaffii*. Regions in gray represent the centromeric sequences amplified in this work. Black boxes indicate identified ARS sequences [11].

LA3 strain was individually transformed with pPICH-ADE3 and centromeric plasmids pPICH-CEN1, 2 and 4, which were verified for autonomous replication by plasmid rescue in *E. coli* following restriction digestion with *Not*I to verify vector integrity (S1 Fig).

Plasmid stability was firstly verified through colony color in non-selective medium (Fig 5). When plated on YPD non-selective medium, colonies transformed with pPICH-ADE3 lost their color rapidly and presented a red center with large white edges, a result consistent with plasmid instability. In contrast, strains transformed with centromeric plasmids presented a uniform red coloration throughout the colony.

**Fig 5.**
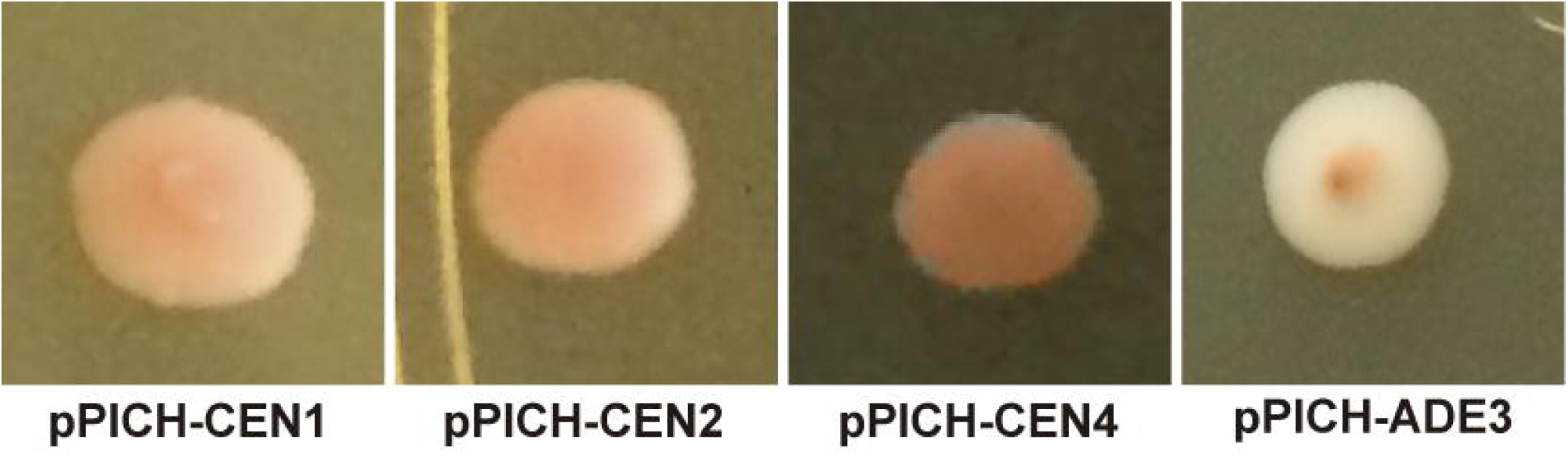
Plasmid stability analysis. LA3 strain transformed with pPICH-ADE3 and the centromeric plasmids was grown on YPD medium for 3 days until color development.

Further stability examination of the centromeric plasmids was performed by growing cells in liquid YPD medium for 144 hours. After diluting and plating cultures on non-selective medium, red and white colonies were counted and compared between each construction (Fig 6). LA3 strain transformed with pPICH-ADE3 did not yield red colonies in any growth period, indicating that the plasmid was mitotically unstable. Conversely, centromeric plasmids presented a higher mitotic stability than pPICH-ADE3. After 96 hours of growth, cells with pPICH-CEN1 started to present white colonies, while the other centromeric plasmids remained stable. After 144 hours, pPICH-CEN1 was lost in most colonies while the other centromeric plasmids were lost in <10% cells. The reason for the instability of pPICH-CEN1 could be related to the absence of an autonomously replicating sequence within the centromere, since centromeres 2 and 4 were cloned with at least partially amplified ARSs. The original replicating sequence in the pPICH-ADE3 plasmid, ARS1, has shown to be less efficient than its modern counterparts, therefore new ARSs contained in the centromeres could have enhanced the centromeric plasmids’ mitotic stability [12].

**Fig 6.**
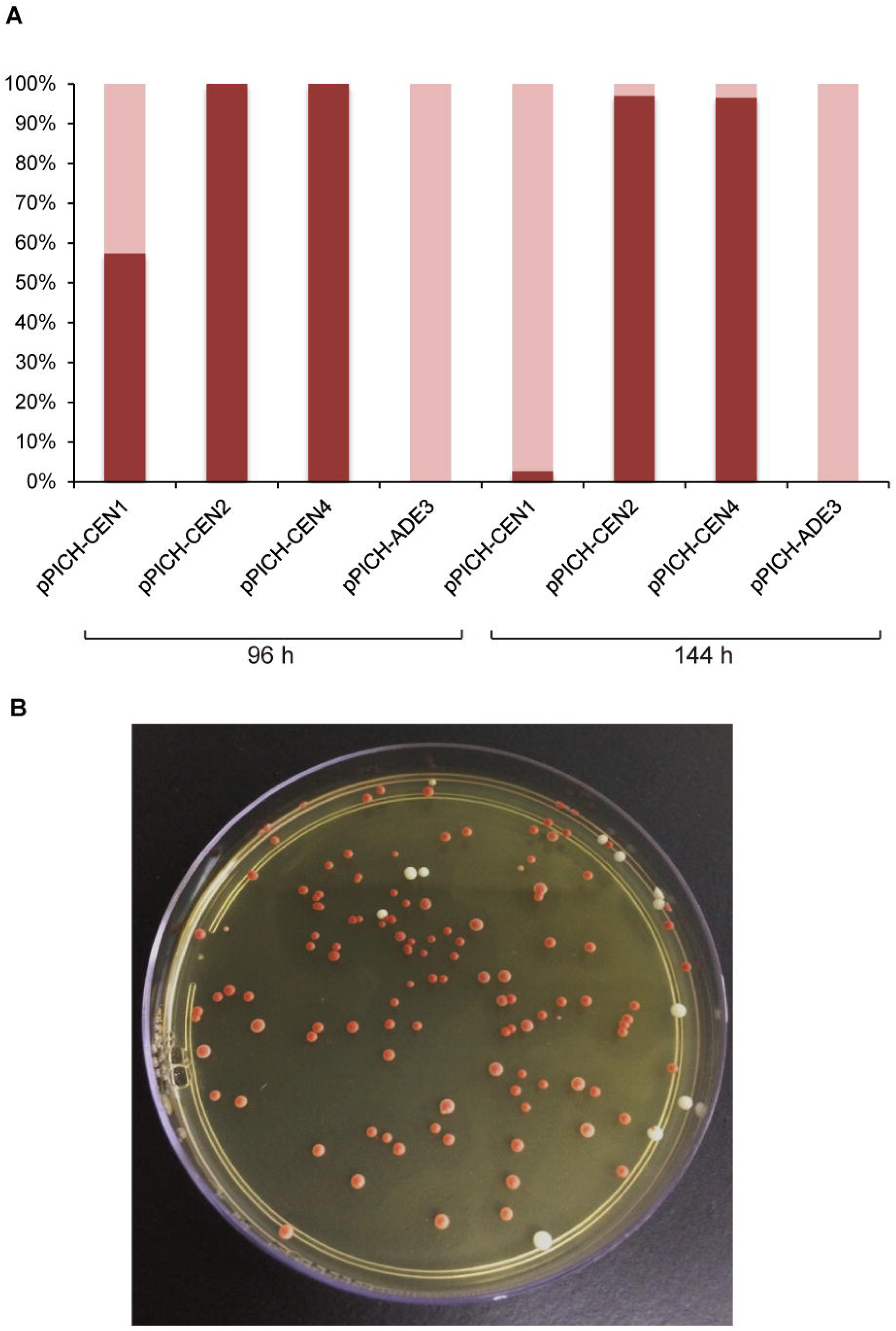
Plasmid stability test. (A) Aliquots of the liquid cultures were collected after 96 and 144 hours of growth and plated on YPD medium. Red colonies represent cells that maintained the *ADE3-*containing plasmid, while white colonies have lost it. Red portions of the bars represent red colonies; light pink bars represent white colonies. (B) A plate representing a typical result after 144 h growth.

Yeast centromeric plasmids knowingly have a higher mitotic stability under non-selective conditions than common replicative vectors since they are equally segregated between daughter cells and therefore provide a uniform culture of cells containing the plasmid [27]. A centromeric vector containing *K. phaffii* CEN2 has been constructed and it presented an enhanced stability when compared to a replicative plasmid [26]. In addition to *K. phaffii* and *S. cerevisiae*, centromeric plasmids have been developed for other yeasts such as *S. pombe, C. glabrata* and *Scheffersomyces stipitis* and in all cases an enhanced plasmid stability under non-selective conditions was verified [13,18].

Yeast replicative plasmids are normally replicated but are unevenly distributed between daughter cells, which creates both multi-copy and plasmidless cells [27]. Under selective conditions, cells lacking the plasmid are unable to survive and the result is a population of multi-copy plasmid-containing cells. The construction of centromeric plasmids should provide better plasmid segregation and stability and cells should maintain a low and stable plasmid copy number during yeast growth [32]. Plasmid copy number was assessed by qPCR after strains were grown in YPD medium containing zeocin in order to ensure that all cells assayed were harboring the centromeric plasmids. The results were compared to the LA3 strain transformed with pPICH-ADE3 also grown in selective medium and to the LA3 control strain, grown in YPD medium. qPCR results (Fig 7) indicate that the strain transformed with centromeric plasmids carried 1-2 copies per cell while the replicative plasmid was present at approximately 25 copies per cell. The difference in plasmid copy number between the replicative vector and the centromeric vectors was significant according to a t-test (p<0.05). This result illustrates the expected segregation pattern described above for growth in selective conditions and, together with the mitotic stability analysis, provides a clear picture of *K. phaffii* genetic manipulation using centromeric plasmids.

**Fig 7.**
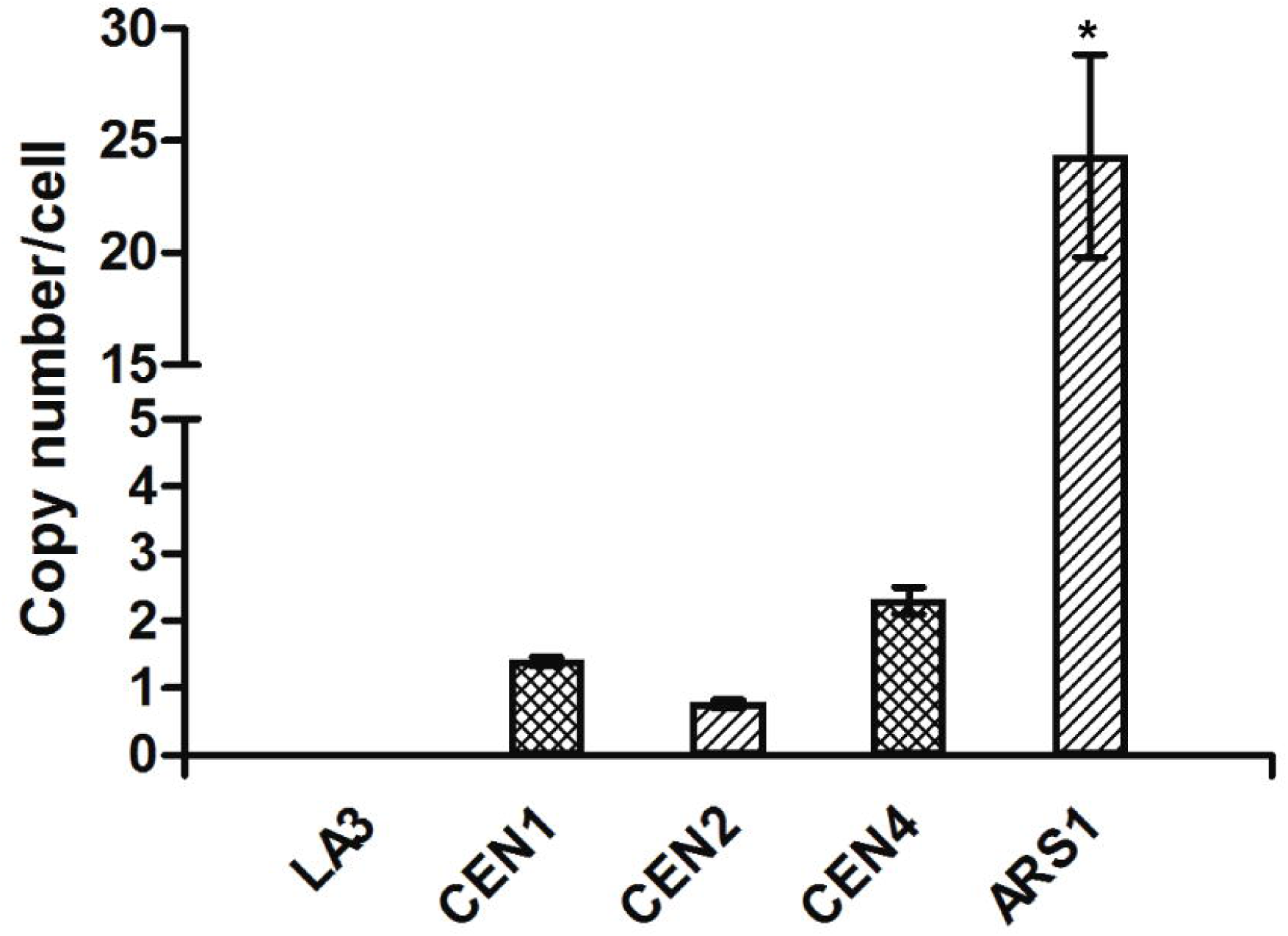
Plasmid copy number determination. The number of plasmids in each cell was estimated by qPCR. LA3 was used as negative control. Statistical analysis comparing each of the centromeric plasmids with the replicative vector was performed through a t-test using GraphPad Prism 5. (p<0.05). Error bars depict the standard deviation of the mean (n = 3).

*S. cerevisiae* centromeric plasmids, in comparison to plasmids bearing the 2 micron sequence, presented the same difference in copy numbers when auxotrophic markers were used. However, when *kanMX* G418 resistance marker was used, the plasmid copy numbers did not differ between centromeric and replicative plasmids [37]. This indicates that factors other than the type of replication origin can influence plasmid copy numbers. In a previous study, *K. phaffii* transformed with a plasmid containing centromere 2 was analyzed regarding plasmid copy number and compared to both a replicative plasmid and an integrative strategy [26]. Results exhibited a low number of plasmids per cell in all strategies, which does not correspond to our observation. The difference may be related to the different strains, culture conditions or plasmid constructions, since although both plasmids used the ARS1 replicative sequence and the zeocin resistance marker, pPICH-CEN1, 2, and 4 contained the *ADE3* gene while the previously reported centromeric plasmid carried an *EGFP* reporter gene.

Overall, our results indicate that centromeric plasmids could be employed as a new tool for the genetic manipulation of *K. phaffii*. Plasmids were maintained for long periods in non-selective medium, indicating that growth can be performed without the addition of antibiotics or any form of selective pressure. The centromeric plasmids’ low copy numbers per cell characterize a stable and homogeneous culture that can provide reliable expression results. Finally, their structure as a circular molecule allows *in vivo* plasmid assembly with relatively short homologous sequences when compared to genomic integration techniques where sequences have to be much longer for directed homologous recombination. Simpler assembly may also facilitate the construction of larger and more sophisticated vectors such as yeast artificial chromosomes (YAC), whose stability features may be also analysed by the color-based assay described in this work.

## Materials and methods

### Strains and Media

DNA cloning was performed using chemically competent *Escherichia coli* XL-10 Gold (Agilent Technologies) grown in LB medium (5 g L^-1^ yeast extract, 10 g L^-1^ peptone and 10 g L^-1^ NaCl, pH 7.2). When needed, agar was added to a final concentration of 1.5%. When zeocin (25 μg mL^-1^) was used for bacterial antibiotic selection, NaCl concentration was reduced to 5 g L^-1^.

*K. phaffii* strains were derived from X-33 (Invitrogen). LA2 strain (*amd2 ade2*) was described in previous work [30]. Yeast was routinely grown in YPD medium (10 g L^-1^ yeast extract, 20 g L^-1^ peptone and 20 g L^-1^ glucose). Solid medium used 2% agar. Zeocin and geneticin, when used, were added at 100 μg mL^-1^ and 500 μg mL^-1^, respectively. Hygromycin B was used to a final concentration of 50 μg mL^-1^. Minimal medium (MD) used 0.34% Yeast Nitrogen Base, 1% (NH_4_)_2_SO_4_, 2% glucose, 0.00004% biotin, and 0.0002% adenine or 0.004% histidine, when needed.

### PCR

DNA was amplified using Invitrogen Platinum Taq DNA Polymerase (High Fidelity), Promega GoTaq Colorless Master Mix or Sigma-Aldrich Accutaq LA DNA Polymerase. All primers used in this work are shown in Table 1.

**Table 1.**
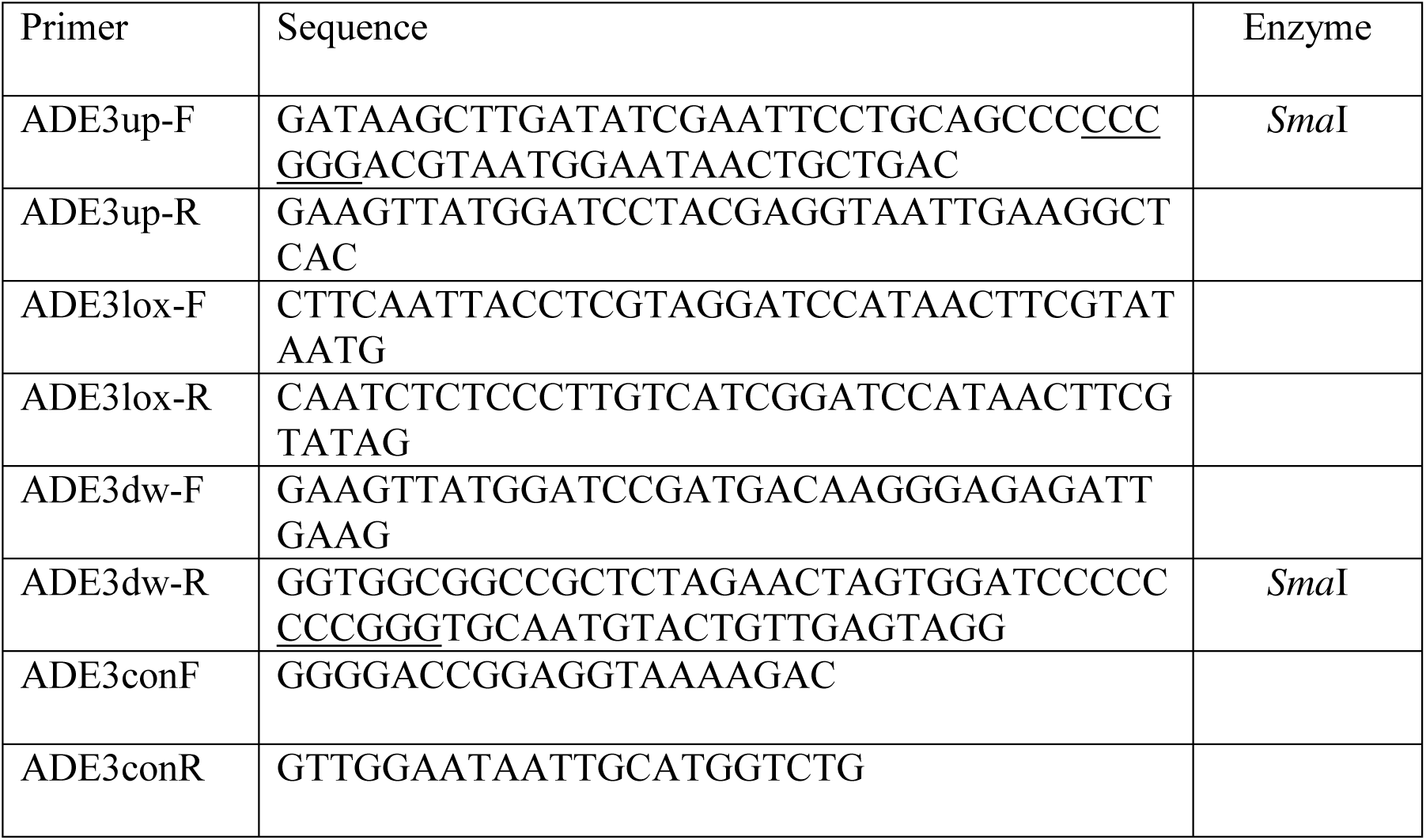
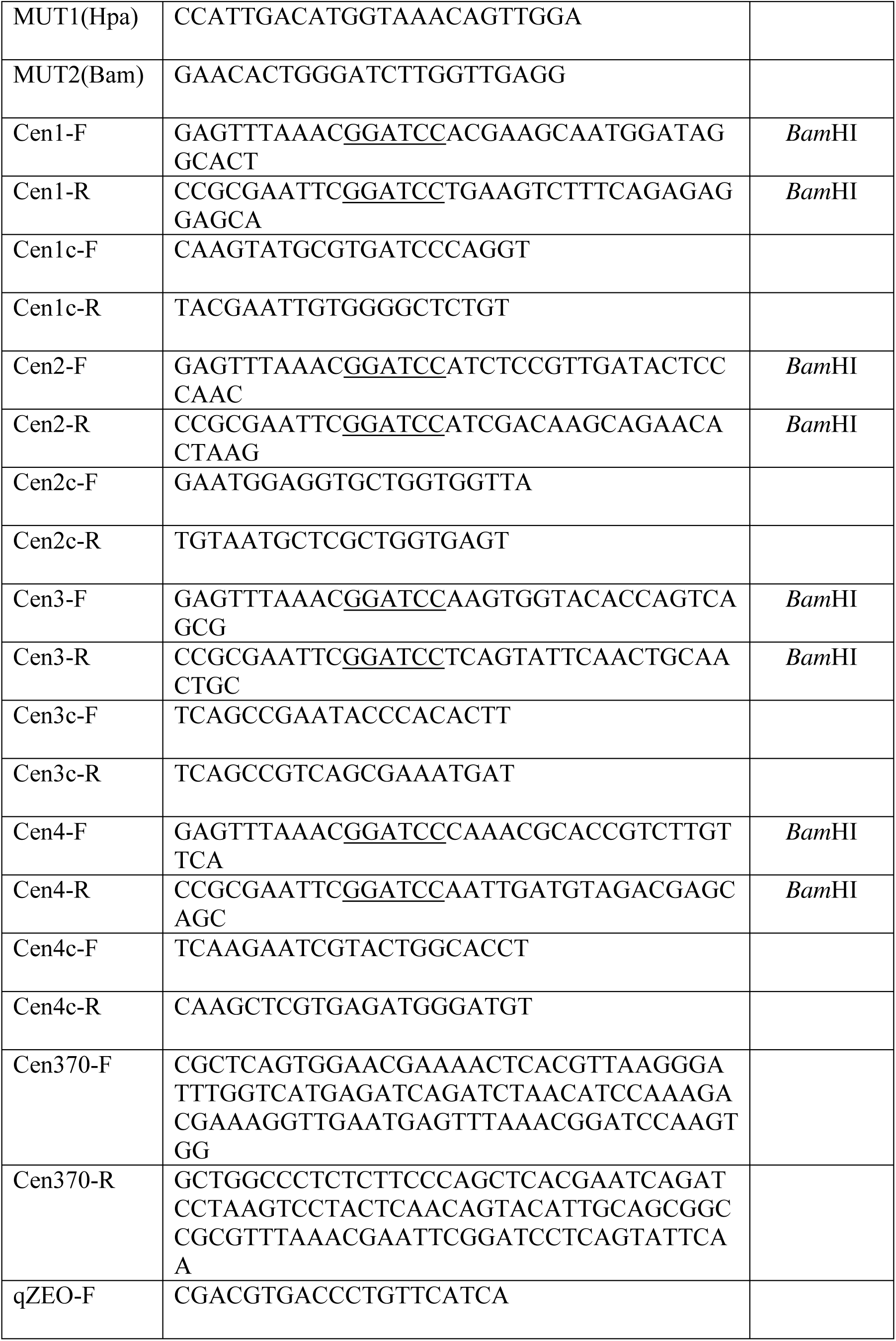
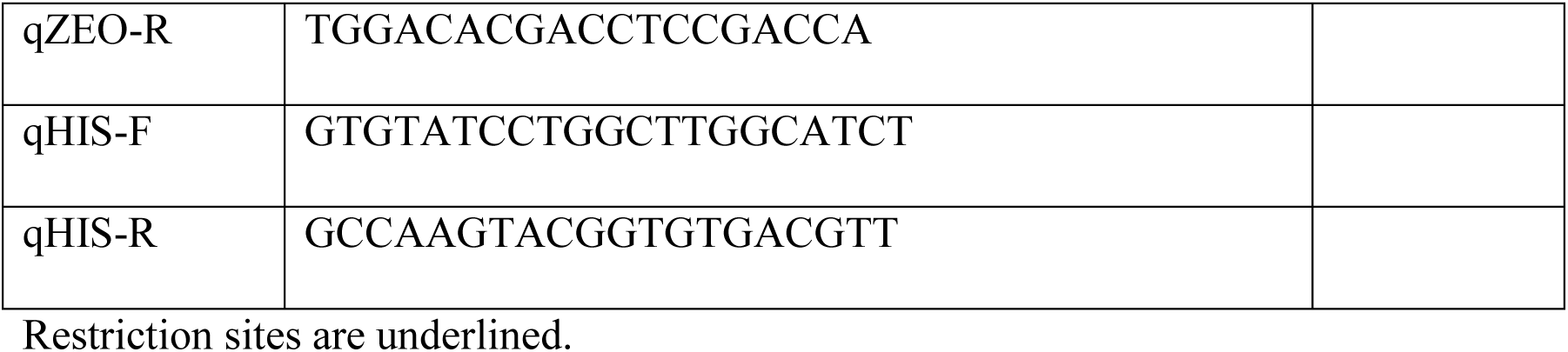
Primers used in this work.

### DNA manipulation

All basic DNA manipulation and analysis were performed as previously described [33]. Restriction digestion was performed in accordance to the manufacturer instructions (New England Biolabs), as well as vector dephosphorylation with Shrimp Alkaline Phosphatase (Promega) and ligation with T4 DNA Ligase (USB). In-Fusion Cloning Kit (Clontech) was used for in-vitro assembly of plasmids. Site-directed mutagenesis was performed using the Transformer Site-Directed Mutagenesis Kit (Clontech). PCR and gel purification used Promega Wizard SV Gel and PCR Clean-Up System.

### Quantitative PCR (qPCR)

Strains harboring the zeocin resistance plasmids were grown to an OD_600_ (optical density measured at 600 nm) of 1 in 10 mL YPD containing zeocin while LA3 was grown in 10 mL YPD. Cells were collected by centrifugation at 2000 x g for 5 minutes. The cell pellet was resuspended with 1 mL 0.25% SDS and incubated at 98°C for 8 minutes according to previous work [26]. Finally, cell debris was removed by centrifugation and DNA was diluted 10-fold in water before qPCR reactions.

Quantitative PCR reactions used primers qZEO-F and qZEO-R for plasmid quantification and qHIS-F and qHIS-R as an internal single-copy control. Assays were carried out with iTaq Universal SYBR Green Supermix (Bio-Rad) in a Rotor-Gene Q (Qiagen) thermal cycler. Analysis used the absolute quantification method and standard curves that ranged from 1×10^4^ to 1×10^8^ copies of the gene of interest. pPIC9 (Invitrogen) and pPICH linearized plasmids were used for construction of the standard curves.

### Yeast transformation

*K. phaffii* was electroporated following two different protocols. For integrative cassettes, we followed the *Pichia* Expression Kit protocol (Invitrogen) and when using replicative plasmids, we proceeded as described previously [34].

### Construction of an *ade2 ade3* strain for color-based stability assays

Strain LA2 [30] was transformed with an *ADE3* deletion cassette and had the marker recycled before transformation with the centromeric plasmids. Construction of the deletion cassette used PCR reactions assembled by an “In-Fusion” cloning reaction. Briefly, primers ADE3up-F and R, ADE3dw-F and R were used for PCR amplification of 491 bp and 582 bp, respectively, from the *K. phaffii* genome. These reactions amplified sequences used for directing homologous recombination and substitution of the complete *ADE3* coding sequence. Meanwhile, primers ADE3lox-F and ADE3lox-R amplified the *kanR* geneticin resistance cassette from plasmid pGKL [35]. PCR fragments were assembled and cloned into pBluescript II SK^+^ linearized with *Sma*I. A final PCR reaction using primers ADE3up-F and ADE3dw-R amplified the whole deletion cassette which was used for transformation of *K. phaffii* LA2. Cells were selected in YPD containing geneticin.

The resulting *ade2 ade3* strain was later transformed with pYRCre2 [36] and selected in YPD supplied with hygromycin B. This step promoted a Cre-mediated excision of the *kanR* cassette eliminating geneticin resistance. After PCR confirmation of marker recycling using primers ADE3conF and ADE3conR, the resulting strain was plated in non-selective YPD medium, causing loss of the pYRCre2 plasmid. The resulting strain was named LA3.

### Construction of centromeric plasmids containing *ADE3*

Plasmid pPICH [30], which is derived from pPICHOLI (MoBiTec), contains the ARS1 replicating sequence [1]. This sequence is originally located on *K. phaffii* GS115 chromosome 2, coordinates 413701-413856 [2]. The plasmid was digested with *Not*I for cloning of the *K. phaffii* native *ADE3*. The complete gene was amplified from X-33 DNA using primers ADE3up-F and ADE3dw-R following digestion with *Not*I. After vector dephosphorylation, fragments were ligated and transformed into *E. coli* XL-10 Gold. One positive clone was then submitted to site-directed mutagenesis using primers Mut1(Hpa) and Mut2(Bam) for removal of the *Bam*HI restriction site present within the *ADE3* coding sequence. The final plasmid containing ARS1, the *Sh ble* resistance marker and *ADE3* was named pPICH-ADE3.

pPICH-ADE3 was digested with *Bam*HI for cloning of all four *K. phaffii* centromeres. These were amplified from *K. phaffii* X-33 genomic DNA using two PCR reactions for each centromeric sequence. Primers Cen1/2/3/4-F and Cen1/2/3/4c-R amplified the first inverted repeat of each centromere while primers Cen1/2/3/4c-F and Cen1/2/3/4-R amplified the other half of the sequences. In order to promote *in vitro*/*in vivo* assembly, the amplicons had ∼80 bp homology between each other in one end and 15 bp with pPICH-ADE3 in the other end. First, we attempted an “In-Fusion” cloning reaction for each of the four centromeres using linearized pPICH-ADE3 and the two PCR fragments. Plasmids were extracted and analyzed by restriction digestion. Centromeres 1, 2 and 4 were successfully assembled and cloned into the plasmid through this strategy. Centromere 3 did not yield any *E. coli* clones following the “In-Fusion” reaction; we then proceeded to an *in vivo* assembly strategy. Primers Cen370-F and Cen3c-R; Cen3c-F and Cen370-R amplified both inverted repeats adding 70 bp of homologous sequences between the fragments and pPICH-ADE3. Finally, we transformed *K. phaffii* LA3 using the linearized vector and both centromeric fragments, using 85 bp of homology for directing recombination. Clones were selected in YPD supplied with zeocin. However, none of the obtained clones presented the assembled plasmid as it was expected and this centromeric sequence was not used in further analyses.

The resulting plasmids were named pPICH-CEN1, pPICH-CEN2, and pPICH-CEN4. All plasmids were transformed into *K. phaffii* LA3 for subsequent stability and quantification assays.

### Stability analysis

LA3 strain transformed with each of the three centromeric plasmids was grown in 20 mL YPD for 16 hours at 28°C and 200 rpm. This culture was inoculated to 20 mL YPD to an initial OD of 0.1. After 24 h of growth under the same conditions the culture was used as inoculum for another flask containing 20 mL YPD to an OD of 0.1. This procedure was repeated every 24 h until a total of 144 hours. At 96 and at 144 hours of growth, a small amount of cells was diluted 10^6^-fold and 100 μL of this dilution were plated on YPD. Plates were incubated at 30 °C for 72 h.

## Supporting information captions

**S1 Fig. Restriction analysis of recovered plasmids pPICH-CEN1, 2, and 4.** Plasmids were digested with *Not*I and analyzed on 1% agarose gel. All digested plasmids yielded a common 4.1 kb band which represents the *ADE3* gene. The size of the upper bands represent the sum of individual centromeres (CEN1 = 7.6 kb; CEN2 = 9.0 kb; CEN4 = 8.7 kb) and other common vector sequences. M: 1 kb Plus DNA Ladder (Thermo Fischer Scientific).

