## Supplementary figures and images for "Cloning and functional characterization of *Komagataella phaffii* centromeres by a color-based plasmid stability assay"

### Supplementary Figure 1

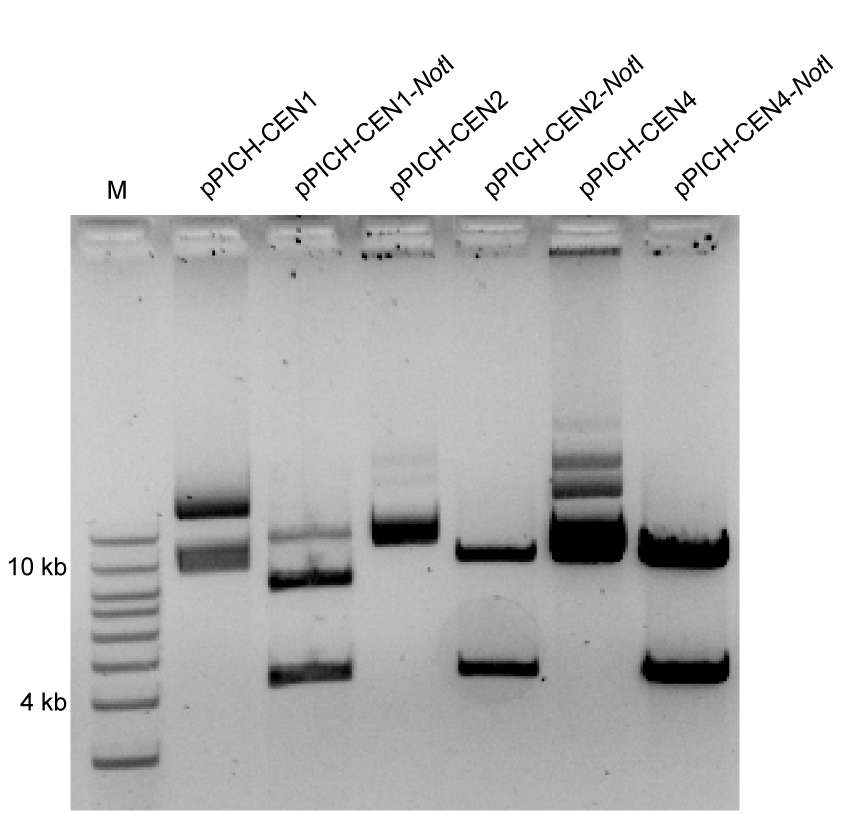
